# An Inexpensive RT-PCR Endpoint Diagnostic Assay for SARS-CoV-2 Using Nested PCR: Direct Assessment of Detection Efficiency of RT-qPCR Tests and Suitability for Surveillance

**DOI:** 10.1101/2020.06.08.139477

**Authors:** Jayeshkumar Narsibhai Davda, Keith Frank, Sivakumar Prakash, Gunjan Purohit, Devi Prasad Vijayashankar, Dhiviya Vedagiri, Karthik Bharadwaj Tallapaka, Krishnan Harinivas Harshan, Archana Bharadwaj Siva, Rakesh Kumar Mishra, Jyotsna Dhawan, Imran Siddiqi

**Author notes:** These authors contributed equally to this work (Jayeshkumar Narsibhai Davda, Keith Frank, Sivakumar Prakash).

## Abstract

With a view to extending testing capabilities for the ongoing SARS-CoV-2 pandemic we have developed a test that lowers cost and does not require real time quantitative reverse transcription polymerase chain reaction (RT-qPCR). We developed a reverse transcription nested PCR endpoint assay (RT-nPCR) and showed that RT-nPCR has comparable performance to the standard RT-qPCR test. In the course of comparing the results of both tests, we found that the standard RT-qPCR test can have low detection efficiency (less than 50%) in a real testing scenario which may be only partly explained by low viral representation in many samples. This finding points to the importance of directly monitoring detection efficiency in test environments. We also suggest measures that would improve detection efficiency.

## Introduction

The continuing Covid-19 pandemic has created an urgent need for increased diagnostic tests worldwide. The requirement of tests has exceeded the normal testing capacities available in public and private hospitals and clinical research laboratories and also strained financial resources. The most widely deployed type of test for identifying individuals infected with SARS-CoV-2, based on recommendations of the World Health Organization (WHO) and national health centres such as the US Centers for Disease Control and Prevention (CDC) detects the presence of viral RNA. The method employs real time quantitative reverse transcription polymerase chain reaction (RT-qPCR) of RNA extracted from nasopharyngeal (NP) swab samples, to measure amplification of a short segment of a viral gene in the course of a PCR reaction following reverse transcription of viral RNA. Performing the RT-qPCR test requires a real time thermal cycler which is an expensive instrument. Most major research laboratories are equipped with only a limited number and smaller laboratories may have none which has placed constraints on the number of tests as well as places for conducting them. Further, the need for fluorescent oligonucleotide probes adds to the cost of the tests. A number of new diagnostic testing methods aimed at reducing the dependence on expensive equipment and kits that are in short supply have been proposed and are under development for deployment (Broughton et al., 2020; Rauch et al., 2020; Yan et al., 2020). However, given the enormous and widespread current need for diagnostic testing in diverse environments it is equally important to increase the utilization of existing research capabilities that have potential but are currently not being used for testing, through the use of tests that employ more widely available equipment and reagents using simple established methods.

In order to extend the scope of diagnostic testing for SARS-CoV-2 we explored a reverse transcription nested PCR (RT-nPCR) approach that does not depend on RT-qPCR but uses standard RT-PCR as part of an endpoint assay. We developed and tested a RT-nPCR protocol comprising a multiplex primary RT-PCR for amplification of four SARS-CoV-2 amplicons and a control human RPP30 amplicon followed by a secondary nested PCR for individual amplicons and visualization by agarose gel electrophoresis. We also examined the use of RT-nPCR in pooled testing and in direct amplification without RNA isolation.

RNA isolated from NP swab samples that had been previously tested using one of two RT-qPCR tests was examined using RT-nPCR and the results compared. We found that taking both standard RT-qPCR tests together, the RT-nPCR test was able to correctly identify 90% of samples detected as positive by RT-qPCR and also detected 13% samples as positive among samples that were negative by the standard RT-qPCR test (likely false negatives). Based on the experimentally measured false negative rate by RT-nPCR tests from this study we estimated that as many as 50% of positive samples may escape detection in single pass testing by RT-qPCR in an actual testing scenario.

## Results

### Nested RT-PCR test development

We designed 40 sets of nested oligonucleotide primers for specifically amplifying different regions of SARS-CoV-2 and not SARS-CoV based on available sequence information (Genbank ID NC_004718.3 for SARS-CoV and MT050493.1 for SARS-CoV-2). These primers were tested for amplification by a primary PCR, followed by a second round of nested PCR. The starting template was cDNA prepared from a pool of RNA isolated from two NP swab samples that had previously been identified as positive using a RT-qPCR diagnostic kit. From a set of 40 candidate amplicons we selected 4 amplicons that gave visible amplification in the primary PCR and strong bands in the secondary PCR reaction as visualized by agarose gel electrophoresis. The primer sequences for these amplicons were checked against 78 SARS-CoV-2 sequences from Indian isolates. Except for a single mismatch with two sequences in the middle for two primers, all primers were 100% identical to all 78 sequences. These amplicons comprised portions of Orf1ab, M, and N genes (Figure 1A; Table 1). The amplified bands were excised from the agarose gel, the DNA was extracted, and identity of each band was confirmed by sequencing. Similarly a set of nested primers was developed for the human RPP30 gene as a control. To test efficiency of amplification we used a dilution series of amplicons as templates in primary and secondary PCR. 1-10 molecules of DNA in dilutions of isolated viral amplicons could be detected by nested PCR (Figure 1B). To detect the presence of SARS-CoV-2 in RNA isolated from NP swabs we performed a multiplex one-step RT-PCR on RNA from positive and negative samples using pooled primers for the four viral amplicons together with human RPP30 control. The product of the primary one-step RT-PCR was used as template in separate nested secondary PCR reactions for each of the amplicons followed by detection using agarose gel electrophoresis. Using the nested RT-PCR assay we were able to detect amplification of the four viral amplicons in RNA from positive samples and no amplification was detected in negative samples (Figure 2).

**Figure 1:**
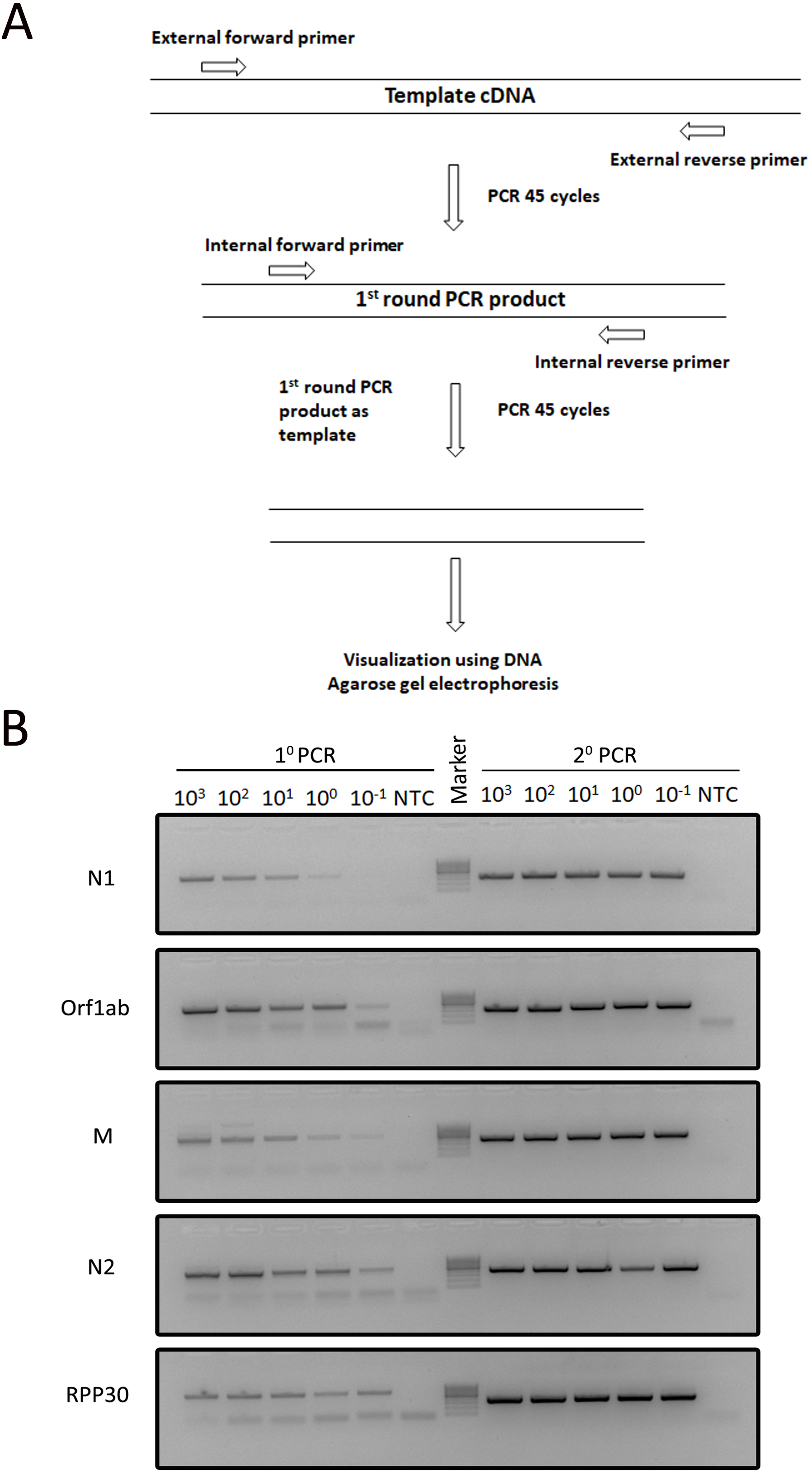
Nested RT-PCR strategy for SARS-CoV-2 detection A) Schematic of nested PCR approach B) Detection limit by serial dilution of amplicons from 1000 – 0.1 molecules.

**Table 1:**
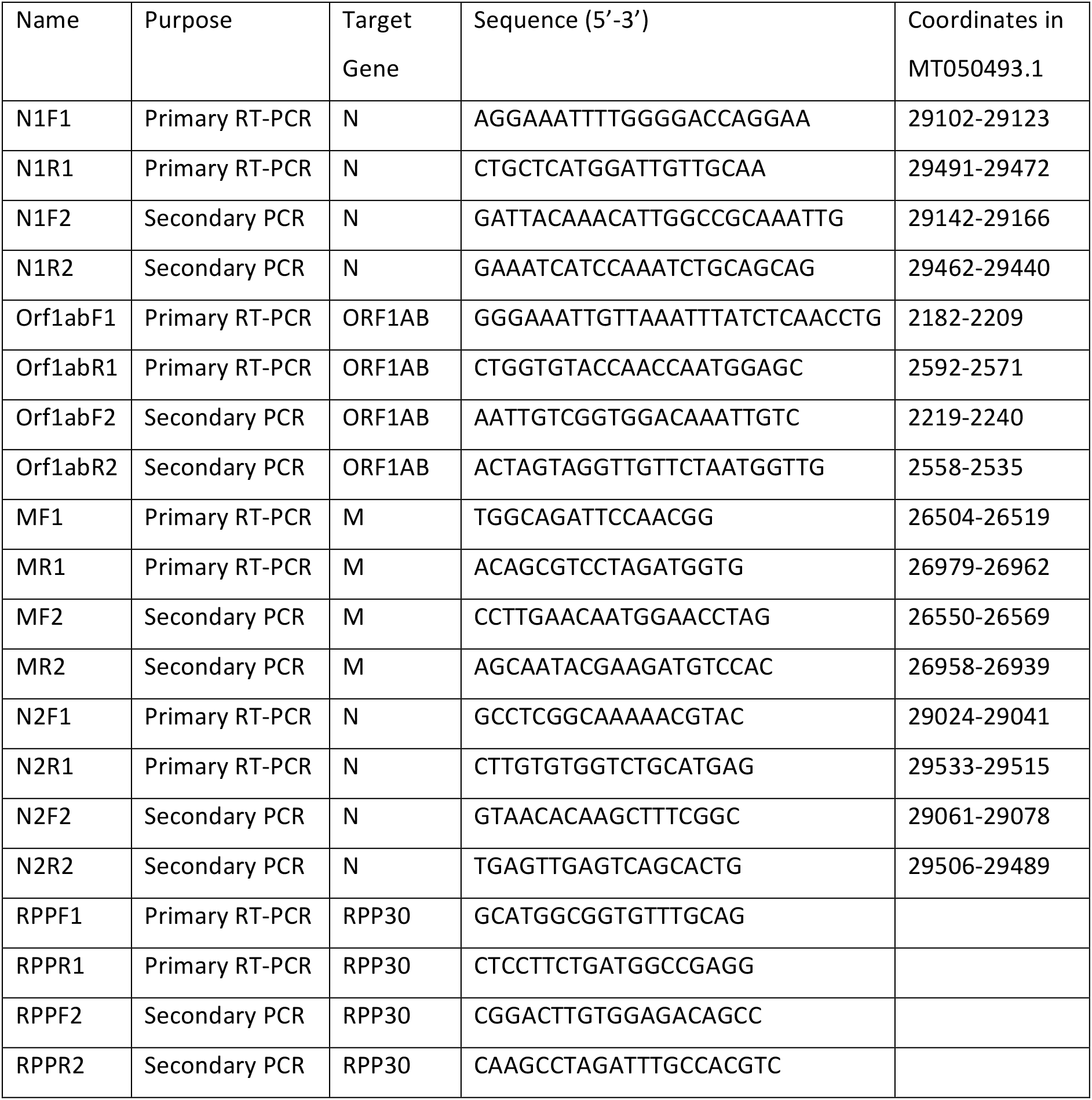
List of primers

**Figure 2:**
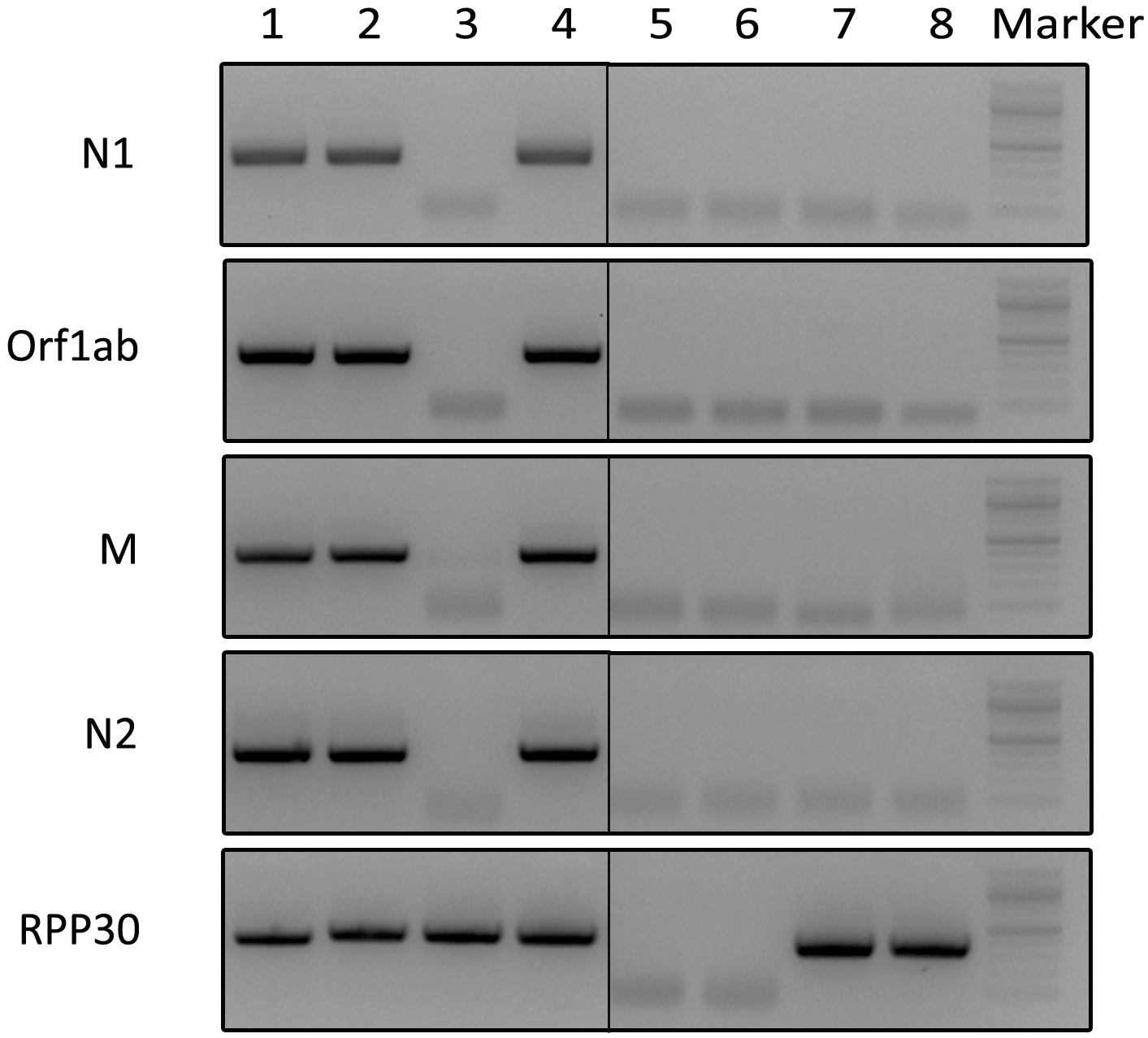
Representative gel picture of positive and negative RNA samples. Lanes 1,2, and 4: positive samples; Lanes 3, 7, and 8: negative samples; Lane 5: 1^0^ NTC; Lane 6: 2^0^ NTC.

### Pooled and direct testing

Pooled testing has been used for SARS-Co-V2 detection (Yelin et al., 2020). In order to assess the suitability of the RT-nPCR test for analysis of pooled samples, we performed the test on three sets of pooled samples comprising RNA isolated from different dilutions of a positive sample in a pool of ten negative samples. For a sample having a cycle threshold (Ct) value of 38.7 for the E gene, the RT-nPCR test gave robust detection in RNA from undiluted sample (positive for 3 out of 4 viral amplicons) and less robustly at a dilution of 1:5 (positive for 1 out of 4 viral amplicons; Figure 3). For two samples having a Ct value of 28.7 and 34.5, the RT-nPCR test successfully detected presence at a dilution of 1:20. Hence the RT-nPCR test was able to detect presence of all three samples in a pool of 1:5.

**Figure 3:**
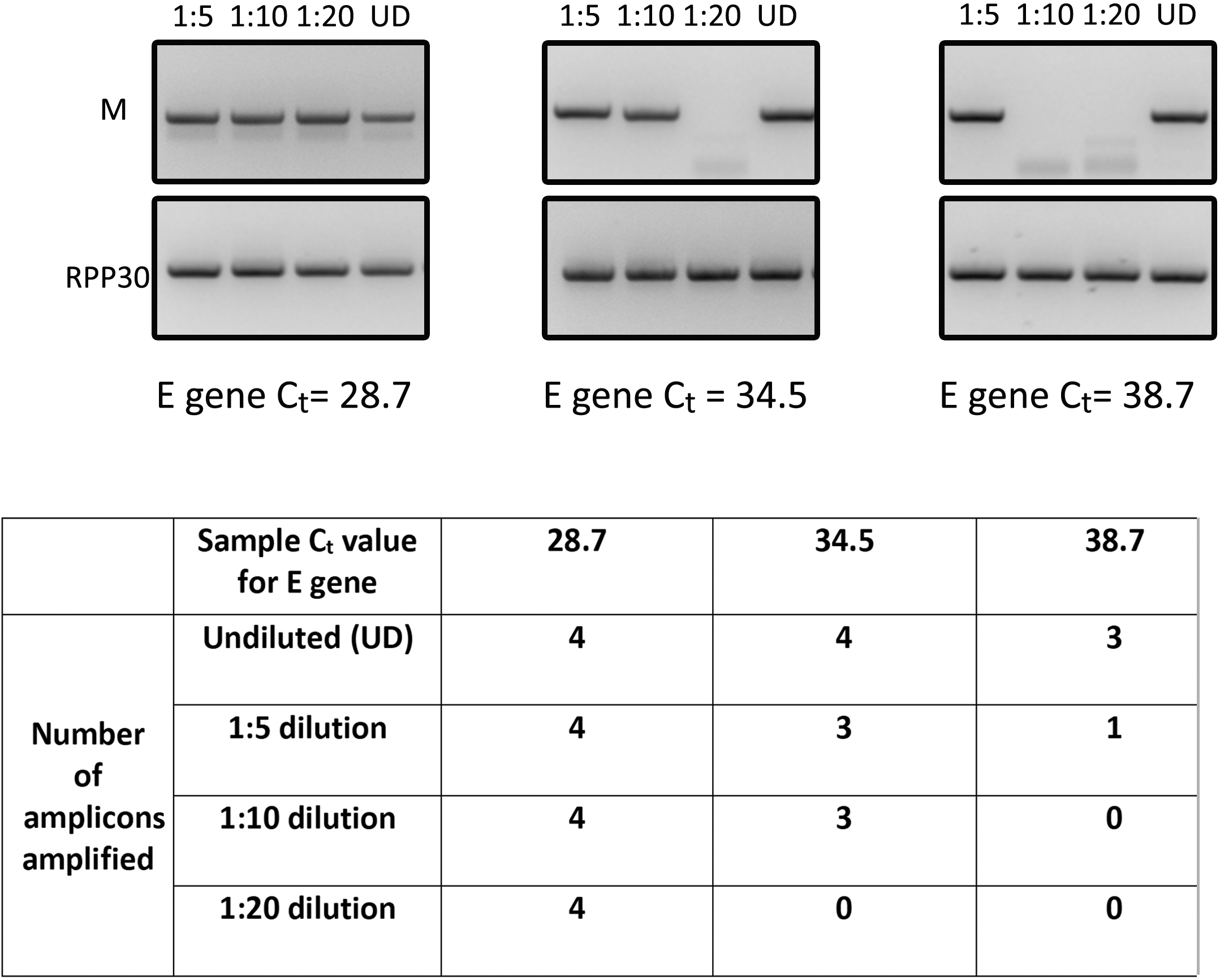
Detection efficiency of RT-nPCR test in pooled samples. Samples with different Ct values for E gene pooled at different dilutions with negative samples. Each pool was tested for amplification of 4 amplicons. UD: undiluted.

Direct testing of samples by RT-qPCR without RNA isolation has been described in recent reports (Bruce et al., 2020; Merindol et al., 2020; Smyrlaki et al., 2020). We performed direct testing of heat inactivated positive samples by the RT-nPCR test. Using 3 μl of swab sample directly, which corresponds to 1/5 of the amount used in testing of isolated RNA, the RT-nPCR test detected a positive sample having an E gene Ct value of 27.9 but not a sample having a higher Ct of 32.7 (Figure 4). Using a higher volume of swab sample gave weaker amplification suggesting the presence of an inhibitor in the VTM. We observed improved amplification when polyvinylpyrrolidone was included in the PCR reaction. Passage of the sample through a Sephadex-G50 spin column also relieved the inhibition and gave improved amplification. Overall the RT-nPCR test is capable of detecting positives directly in swab samples but at a reduced sensitivity compared to isolated RNA.

**Figure 4:**
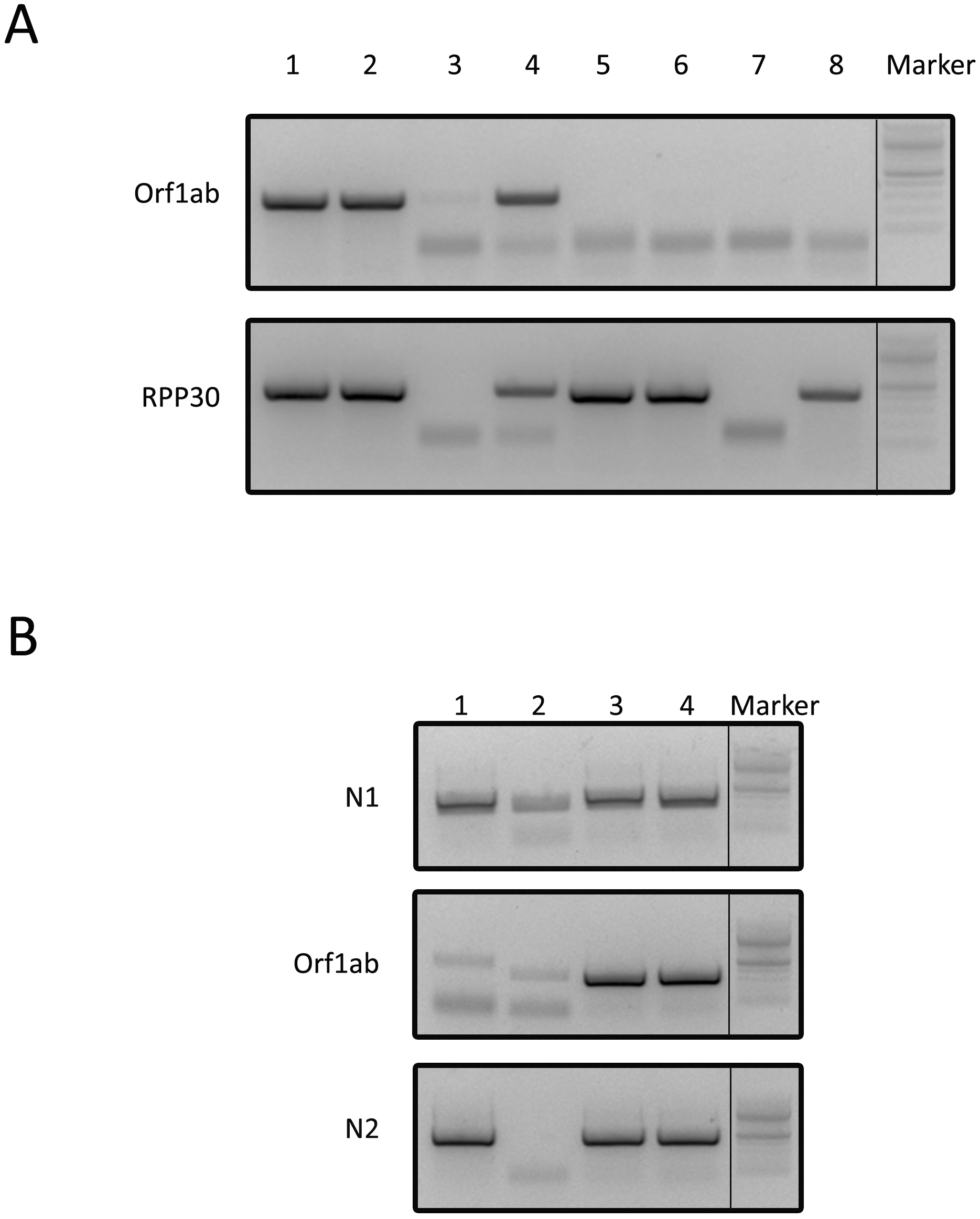
Detection of SARS-CoV-2 by RT-nPCR without RNA isolation. A) Relief of inhibition at higher sample volumes by adding 0.5 % of PVP. Lanes 1-4: E gene Ct = 27.9; Lanes 5-8: E gene Ct = 32.7. Lanes 1,5: 1 μl; 2,6: 3 μl; 3,7: 5 μl; 4,8: 5 μl + 0.5 % PVP. B) Relief of inhibition at higher volume of NP swab sample by passing through Sephadex G-50 spin column. Lane 1, 2: 3 μl and 9 μl of NP swab sample. Lane 3, 4: 3 μl and 9 μl of NP swab sample after passing through Sephadex G-50 column.

### Comparison of RT-nPCR with RT-qPCR

In order to compare the performance of RT-nPCR with that of standard RT-qPCR we tested RNA samples that had tested positive by RT-qPCR. Samples that were positive for at least two amplicons out of four by RT-nPCR were called positive, samples that were positive for one amplicon were considered ambiguous and not included in the comparison, and samples that were negative for all four amplicons were called negative. We compared RT-nPCR test results to sets of positive and negative RNA samples that had been tested by either of two RT-qPCR protocols, one provided by National Institute of Virology (NIV), India (derived from Charite, Berlin and CDC, USA protocols), and the other a commercial LabGun RT-qPCR kit (LabGenomics, Korea). The results indicated that the agreement between the RT-nPCR and NIV tests could be rated as moderate (Cohen’s Kappa = 0.612, 95% CI: 0.42-0.81) (McHugh, 2012) while that between the RT-nPCR and LabGun tests could be rated as almost perfect (Cohen’s Kappa = 0.905, 95% CI: 0.80-1.0;)(Figure 5). The lower agreement with the NIV set was due to 10 samples that were negative by the NIV protocol but positive in the RT-nPCR test. The mean number of positive amplicons for these 10 samples was 3.3 indicating that they were likely to be true positives (i.e. false negatives). Based on the number of RNA samples that were negative in the RT-qPCR tests but positive by RT-nPCR we derived an estimate of the proportion of positive samples that were being detected by the NIV and LabGun RT-qPCR tests (Table 2). The NIV set covered 400 samples and had a prevalence rate (% positives) of 15%. The detection efficiency for this set was calculated to be 0.47. The LabGun set comprised 1186 samples and had a prevalence rate of 3.7%. The detection efficiency for the second set was estimated to be 0.44. Therefore our estimate suggests that a high proportion of positive samples (upto 50% or more) can be missed by the standard RT-qPCR test in a real testing scenario.

**Figure 5:**
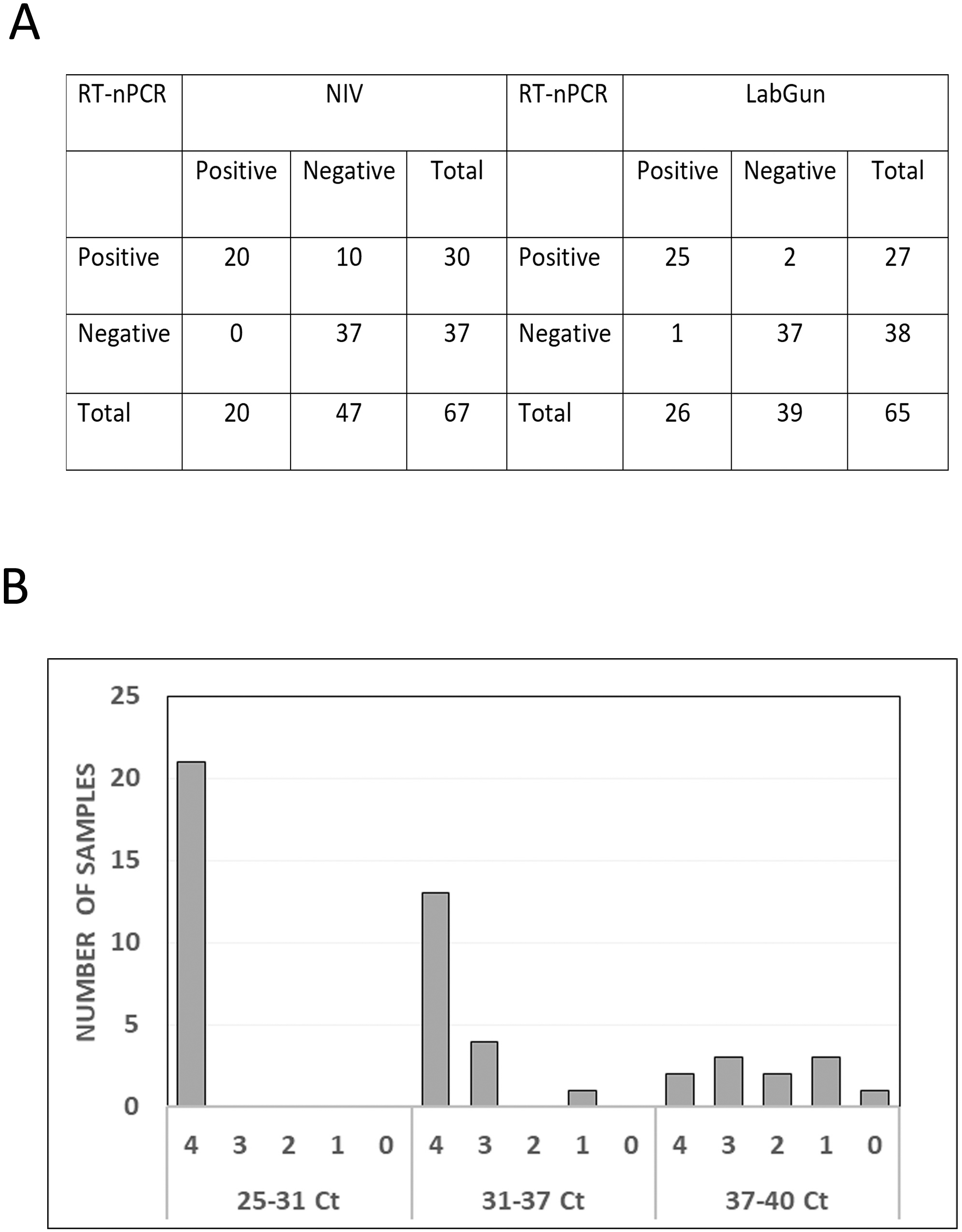
Comparison of RT-nPCR and RT-qPCR tests A) Contingency table of RT-nPCR with RT-qPCR tests. Samples that were positive for only one amplicon by RT-nPCR were considered ambiguous and excluded from analysis. The number of ambiguous samples was: from NIV set: Positives 0/20, Negatives 3/50; LabGun set: Positives 4/30, Negatives 2/41. B) Graph showing number of amplicons amplified by RT-nPCR of samples having a range of Ct values for E gene. Ct 25-31 (n = 21), Mean = 4.0; Ct 31-37 (n = 18), Mean = 3.61; Ct 37-40 (n = 11), Mean = 2.18.

**Table 2:**
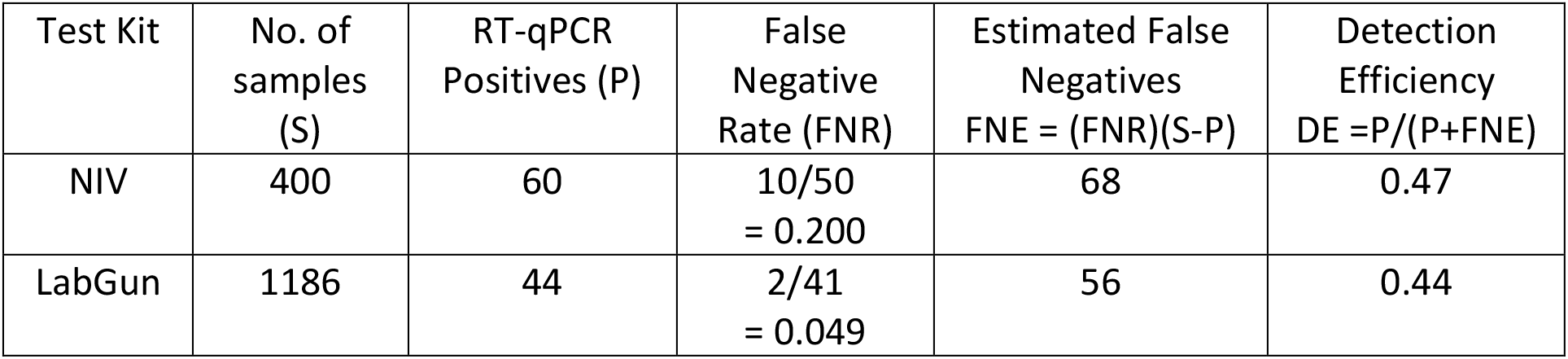
Estimate of Detection Efficiency of Positives by RT-qPCR

| Test Kit | No. of samples (S) | RT-qPCR Positives (P) | False Negative Rate (FNR) | Estimated False Negatives<br>$FNE = (FNR)(S-P)$ | Detection Efficiency<br>$DE = P/(P+FNE)$ |
| --- | --- | --- | --- | --- | --- |
| NIV | 400 | 60 | $10/50$<br>$= 0.200$ | 68 | 0.47 |
| LabGun | 1186 | 44 | $2/41$<br>$= 0.049$ | 56 | 0.44 |

## Discussion

The RT-nPCR test described above does not require a real time thermal cycler and can be performed in a laboratory that has basic molecular biology equipment, a thermal cycler, and a BSL2 room with Class II laminar flow hood. It uses well-established methodologies and reagents that are widely available and can serve as a basis for broadening the scope of testing for SARS-CoV-2. The performance of the RT-nPCR test was comparable to RT-qPCR and the cost of consumables for the test based on list prices is around US $ 7 per test about half of which is the cost of RNA isolation. The test uses two rounds of PCR amplification which does increase potential for contamination. However we found that by following a set of practices described in a detailed protocol (Supplementary file 1), contamination could be avoided.

In the course of comparing the RT-nPCR and standard RT-qPCR tests we estimated from the rate of experimentally determined false negatives, that a high proportion (about 50%) of positive samples were being missed by the RT-qPCR test applied in a single pass testing protocol. This is a high escape rate which poses a concern in individual diagnosis and merits monitoring and greater comparison of results across testing scenarios. We also suggest that detection can be improved by repeat testing of isolated RNA and by increasing the number of amplicons tested.

The existence of false negatives in the RT-qPCR test has been inferred in a number of studies that compared clinical test and symptomatic data with RT-qPCR testing information (Kucirka et al., 2020; Li et al., 2020; Xiao et al., 2020). Several studies have compared the performance of different RT-qPCR kits on RNA from clinical samples primarily to assess the performance of the kits, and have found agreement as well as differences that also point to the existence of false negative test results (Hogan et al., 2020; Pujadas et al., 2020; van Kasteren et al., 2020; Xiong et al., 2020). However, direct experimental analysis of RT-qPCR negative RNA samples at testing centres using different but related RT-PCR based tests as a way of estimating detection efficiency has received limited attention likely because of the high demand for diagnostic tests, shortage of testing kits, and cost of testing. The low detection efficiency that we have estimated may not be entirely explained by a low representation of the virus in many of the samples as the mean number of positive amplicons by RT-nPCR for the 12 false negatives (10 NIV + 2 LabGun) was 3.25 (Figure 5). Possible alternative explanations are variability in test performance or in representation of different portions of the viral RNA. Low detection efficiency could be a possible concern in other tests including those under development that are based on reverse transcription as a first step followed by DNA amplification and detection (Broughton et al., 2020; Rauch et al., 2020; Yan et al., 2020). Many RT-qPCR tests use RNA corresponding to about 1% of swab sample. Increasing the amount of sample for RNA isolation and concentration of the sample are possible options for improving detection, however these would also increase the number of operations and expense, and it would need to be assessed whether this would be compatible with medium to high throughput protocols. Protocols based on initial detection of a single amplicon followed by a confirmatory test for a second amplicon may also contribute to false negatives and lower detection.

In conclusion the RT-nPCR endpoint assay for SARS-CoV-2 described above uses widely available reagents, lowers costs, and obviates the need for a real time time thermal cycler. The assay can therefore be performed in a large number of clinical and diagnostic laboratories lacking this expensive piece of equipment essential for RT-qPCR testing. The performance of RT-nPCR is comparable to RT-qPCR and analysis of RT-qPCR tested samples by RT-nPCR shows a high escape rate in detection of positives, highlighting a need for directly monitoring detection efficiency through assessment of false negatives in real testing environments. As more regions of the world move into community transmission phase, there is also a greater need for surveillance testing. Pooled testing by RT-nPCR can contribute to meeting this requirement.

## Materials and Methods

### Sample Collection

NP swab samples were collected from patients suspected of being infected with SARS-CoV-2 and their contacts at different hospitals in the Hyderabad vicinity based on Indian Council of Medical Research (ICMR) guidelines (http://www.nie.gov.in/images/leftcontent_attach/COVID-SARI_Sample_collection_SOP_255.pdf) and in accordance with Institutional Ethics Committee guidelines. Samples were coded and anonymized before processing and data collection.

### RNA isolation

140 μl of NP swab sample in Viral Transport Medium (VTM) was used for RNA isolation by the QIAamp^®^ Viral RNA (cat# 52906) or equivalent kit as per manufacturer’s instructions. NP sample was lysed in BSL3 facility. RNA isolation steps were carried out in BSL2 facility. RNA was eluted in 50 μl of water. For pooled sample RNA isolation, positive sample was pooled with negative samples in 1:5, 1:10 and 1:20 ratios comprising 140 μl of NP sample and isolated as above. All safety precautions were followed as per ICMR-NIV guidelines.

### Primer design and PCR

SARS-CoV-2 full length genomic sequence of an isolate from Kerala - India was downloaded from NCBI (Genbank ID MT050493.1). Primers were designed for regions specific for SARS-CoV-2, but not SARS-CoV (Genbank ID NC_004718. For the human RPP30 gene (Genbank ID U77665.1) primers were designed on exon-exon junctions to avoid genomic DNA amplification. Oligonucleotides were obtained from Bioserve, Hyderabad. Pooled RNA from two previously identified positive NP swab samples was used for first strand cDNA synthesis (Takara primescript kit cat # 6110A). 20 μl of the cDNA reaction was diluted ten-fold and used for primary PCR. Primary PCR was performed using EmeraldAmp^®^ GT PCR Master Mix (Takara cat# RR310A) 10 ul, forward and reverse primers (5 μM) 1 μl each, diluted cDNA 3 μl, and water 5 μl. Thermal cycling conditions were 1) 95 °C - 2 minutes 2) 95 °C - 15 seconds 3) 60 °C - 15 seconds 4) 72 °C - 30 seconds 5) Repeat steps 2-4 for 4 cycles 6) 95 °C - 15 seconds 7) 55 °C - 15 seconds 8) 72 °C - 30 seconds 9) Repeat steps 6-8 for 39 cycles 10) 72 °C - 2 minutes 11) 15 °C – hold. The primary PCR product was diluted fifty-fold and 1 μl was used for secondary PCR using a nested primer pair. Thermal cycling conditions were the same as for primary PCR. PCR products were separated by electrophoresis on a 1.5% agarose gel and visualized on a UV gel documentation system. For amplicon dilution experiments, bands were excised from the gel, DNA was extracted (Macherey-Nagel Nucleospin kit, Cat # 740609.250) and subcloned into pGEM-T vector (Promega). The insert was PCR amplified from a plasmid DNA clone, followed by gel purification and extraction. DNA was quantified by a Qubit dsDNA HS kit (Thermofisher Cat # Q32851).

### RT-nPCR

Primary RT-PCR and secondary PCR was performed using the Primescript III 1-step RT-PCR kit and EmeraldAmp GT PCR Mix (Takara; detailed protocol provided in Supplementary File 1). For direct testing of sample without RNA isolation NP sample was heat inactivated at 95 °C for 10 minutes in BSL3. 1 μl of 5 % PVP40 (Sigma cat# PVP40-500G) was included in Primary PCR and other conditions were kept same. For Sephadex G50 (Sigma cat# GE17-0042-01) experiment, a spin column was prepared as mentioned in Sambrook and Maniatis manual and other conditions were kept same. The spin column was used outside the PCR area to avoid contamination from aerosols. Images were edited using Adobe Photoshop and included removal of intervening lanes in the gel between samples and DNA marker indicated by a vertical line.

### RT-qPCR

#### NIV method

RT-qPCR for E gene and RPP30 gene was done with 5 μl of RNA template by following First line screening assay according to National Insititute of Virology (ICMR-NIV) https://www.icmr.gov.in/pdf/covid/labs/1_SOP_for_First_Line_Screening_Assay_for_2019_nCoV.pdf. Based on E gene result RdRp and Orf1b were tested with 5 μl of RNA sample by following Confirmatory assay given by ICMR-NIV https://main.icmr.nic.in/sites/default/files/upload_documents/2_SOP_for_Confirmatory_As_say_for_2019_nCoV.pdf.

#### LabGun method

RT-qPCR for E gene and RdRp was performed along with internal control as per manufacturer’s instructions (LabGenomics - LabGun™ COVID-19 RT-PCR Kit cat# CV9032B). 4 μl of RNA template was used for the reaction.

## Ethical statement

The work described in this study was carried out in accordance with institutional ethics committee guidelines.

## Author contributions

JND, KF, SP, and IS designed the experiments. JND, KF, SP, GP, DPV, and DV performed experiments. KBT, KHH, RKM, and ABS supervised processing and provision of samples and data. JND, KF, SP, IS and JD interpreted the experimental results. IS, JND, SP, and KF wrote the paper with inputs from JD.

## Acknowledgements

The Telengana state government and Directorate of Medical Education are acknowledged for providing the swab samples used in this study. We thank the COVID-19 team of volunteers at CCMB for processing the samples. The full list of volunteers is provided as Supplementary file 2. This work was supported by the Council of Scientific and Industrial Research (CSIR, India).

