## Supplementary File 1 for "An Inexpensive RT-PCR Endpoint Diagnostic Assay for SARS-CoV-2 Using Nested PCR: Direct Assessment of Detection Efficiency of RT-qPCR Tests and Suitability for Surveillance"

### Standard Operating Procedures for RT-nPCR Test for SARS-CoV-2 Detection

This protocol assumes that the practitioner has training in molecular biology and is experienced in methods for isolation, processing, and analysis of RNA and DNA.

#### Avoiding carryover contamination:

The Nested RT-PCR Test for SARS-CoV-2 detection provides a high level of sensitivity. In order to reliably perform this test in an environment which involves repeated handling of reagents and analysis of a large number of samples it is critical to avoid carryover contamination of reactions, reagents, and templates while performing the test. Since the Nested RT-PCR Test involves successive rounds of PCR amplification it is extremely sensitive to contamination in even a small amount. It is very easy for contamination to arise and spread across the working environment through aerosols. We provide a detailed protocol below covering conditions which in our experience provided for stable maintenance of contamination free performance over repeated rounds of testing. Ideally 2 people should work together with one person preparing the mixes and the other person bringing and returning the reagents to avoid carryover contamination of PCR reagents and primers.

#### Workspace requirements:

- Pre-PCR:

  Aerosols generated in handling of PCR products readily contaminate workspace and affect results, particularly for nested PCR. We therefore used separate rooms for assembling primary and secondary PCR reactions, each room being equipped with a vertical laminar flow Class II biosafety cabinet having UV lights. The rooms had overhead UV lights operated from an external switch. The rooms were separate from and did not have common air-conditioning with other rooms where PCRs and gels were run.

- PCR:

  Thermal cyclers were placed in a room that was well isolated from the PCR assembly rooms

- Gel Electrophoresis:

  Agarose gel electrophoresis was done in a separate room from the thermal cycler rooms and the biosafety cabinet rooms

#### Pipetmen and Plasticware:

- Pipetmen were decontaminated at the start of the project by disassembling and soaking the barrel and moving parts inside in 0.1N HCl for 15 min, then in 0.1% bleach, then extensive washing in tap water, and finally soaking in MQ water for 15 min followed by air drying. The decontamination was done in a room that was not used for PCR and running gels.
- Pipetmen sets used for RT-PCR and PCR reagents and primers were strictly separate from the set used for handling template. Care was taken to ensure that the Pipetmen used for PCR reagents and primers were never used for template or the product of 1°

PCR. Also the Pipetman used for adding the 1° PCR reaction product to the 2° PCR reaction was exclusive and separate from the one used for adding template to the 1° PCR reaction. These Pipetmen sets were labelled R (Reagents), T1 (Template PCR1), T2 (Template PCR2). These pipettes were not taken into rooms where thermal cyclers and gels were run. A different set of pipettes were used for loading gels.

- Plasticware used for PCR reactions was stored in a separate room from the ones used for thermal cyclers and running gels and kept in secondary plastic containers. Only sterilized aerosol resistant filter tips were used for handling RTPCR/PCR reagents and ALL steps in setting up reactions.

Handling Reagents and Plasticware:

RT-PCR Mix, PCR Mix, primers (5 µM), and 1° PCR primer pool (4 µM each) were stored in aliquots. The protocol below separated the handling of PCR reagents from subsequent template handling steps. Powder-free nitrile gloves were used when handling PCR reagents. When adding template in the 1° PCR, template tubes were opened and closed one at a time before and after each template addition. Care was taken to avoid contact with inside surfaces of tube. Opening of template tubes was done using a folded tissue paper to hold the top of the tube and the paper was discarded after opening. Following addition of all templates the gloves were decontaminated (with 0.1% bleach and 70% ethanol).

Reagents**:**

Primescript III 1-step RT-PCR kit (Takara cat no. RR600A)
EmeraldAmp GT PCR Master Mix (Takara cat no. RR310A)
Oligonucleotide primers (see Table 1 in main text)

**Primary 1-step RT-PCR**

Preparation of Master and Primer Mixes (in biosafety cabinet)

1. Turn on UV in biosafety cabinet 1 and Room 1 for 30 min. Turn off UV. Wipe inside surface of biosafety cabinet with 0.1% bleach followed by 70% Ethanol. Wipe Reagent Pipetmen (R) and Template Pipetmen (T1) with 0.1% bleach followed by 70% EtOH. Place in biosafety cabinet.

2. Bring an aliquot of the Takara Primescript III 2X reaction mix and 1° PCR primer pool (4 µM each primer) from -20 °C freezer (store RT-PCR reagents and primers in a distant freezer from the template RNA (kept at -80 °C)).

3. Using Reagent Pipetmen assemble in the given order:
   Master Mix I:
   Takara Primescript III 2x Reaction Mix 10.0 µl x (n+1 including no template control -- NTC ).
   Close the Master Mix 1 tube.

Primer Mix (minus RNA):

| | |
|---|---|
| 1° PCR primer pool (4 uM each) | 1 µl x (n+1) |
| MQ-water | 4 µl x (n+1) |

4. Close 1° Primer Pool and Primer Mix tubes. Return Primescript III and 1° Primer Pool tubes to -20 °C freezer.

Primer Annealing:

5. Bring RNA template tubes into biosafety cabinet.

6. Aliquot 5 µl Primer Mix (minus RNA) into 'n' PCR plate wells.

7. Change Pipetman to Template P20 (P20T1) and add 5 µl RNA to corresponding wells. Seal wells.

8. Heat at 70 °C for 5 minutes on thermal cycler. Chill on ice. Spin down for 5 seconds (This last step is done outside the biosafety cabinet).

Master Mix 1 Addition:

9. Add 10 µl of Master Mix 1 into respective wells containing RNA+primer using Template P20 (P20T1) or a dedicated multichannel pipetman (NOT Reagent Pipetman). Mix by slow pipetting up and down 2-3 times avoiding air bubbles. Seal plate and run RT-PCR:

| | | |
|---|---|---|
| 52 °C | 30 minutes | (preset at 52 °C before placing plate) |
| 95 °C | 2 minutes | |
| 95 °C | 15 secs | 5 cycles |
| 60 °C | 30 secs | |
| 68 °C | 30 secs | |
| 95 °C | 15 secs | 40 cycles |
| 55 °C | 30 secs | |
| 68 °C | 30 secs | |
| 68 °C | 1 min | |
| 15 °C | forever | |

Return RNA to -80 °C freezer.

Wipe Pipetmen with 0.1% bleach and 70% Ethanol.

Clear and seal waste and clean biosafety cabinet. Turn on biosafety cabinet UV and room UV for 30 minutes.

**Secondary PCR**

Master Mix Preparation:

1. Turn on UV in biosafety cabinet 1,2 and Room 1,2 for 30-60 min. Turn off room UV immediately before re-entering that room and biosafety cabinet UV before use.

2. Bring Emerald 2X Master Mix aliquot to biosafety cabinet 1 and prepare 2° Mixes 1-5 using Reagent Pipetman:
   First add 10 ul x (n+1) 2X Emerald MM into each of 5 tubes: 2°M1-2°M5
   Close 2X Emerald MM and return to 4 °C (short term use) or -20 °C (long term).

3. Bring 2° primers and add to respective 2°M1-2°M5 tubes:

| | | | |
|---|---|---|---|
| 2°M1: | 2X Emerald | | 10 ul x (n+1) |
| | Primer N1F2 | (5 µM) | 1 ul x (n+1) |
| | Primer N1R2 | (5 µM) | 1 ul x (n+1) |
| | MQ water | | 7 ul x (n+1) |
| 2°M2: | 2X Emerald | | 10 ul x (n+1) |
| | Primer Orf1abF2 | (5 µM) | 1 ul x (n+1) |
| | Primer Orf1abR2 | (5 µM) | 1 ul x (n+1) |
| | MQ water | | 7 ul x (n+1) |
| 2°M3: | 2X Emerald | | 10 ul x (n+1) |
| | Primer 67_MI(FP) | (5 µM) | 1 ul x (n+1) |
| | Primer 68_MI(RP) | (5 µM) | 1 ul x (n+1) |
| | MQ water | | 7 ul x (n+1) |
| 2°M4: | 2X Emerald | | 10 ul x (n+1) |
| | Primer MF2 | (5 µM) | 1 ul x (n+1) |
| | Primer MF2 | (5 µM) | 1 ul x (n+1) |
| | MQ water | | 7 ul x (n+1) |
| 2°M5: | 2X Emerald | | 10 ul x (n+1) |
| | Primer N2F2 | (5 µM) | 1 ul x (n+1) |
| | Primer N2F2 | (5 µM) | 1 ul x (n+1) |
| | MQ water | | 7 ul x (n+1) |

Return 2° primer tubes to -20 °C freezer.

Aliquot 19 µl of above mixes into respective wells ( 5 wells total per sample) and seal 2° PCR plate. Place on ice.

Template Addition:

Collect 1° PCR plate from thermal cycler Wipe outside with 0.1% bleach and 70% EtOH. Place on ice for 1 min. Spin 20 secs to pellet liquid to bottom of wells. Take to biosafety cabinet 2.

4. Wipe biosafety cabinet 2 and 1° PCR plate again with 0.1% bleach and 70% EtOH.

5. Carefully open 1° PCR plate/tube in biosafety cabinet using tissue paper covering to reduce spread of aerosols and dilute 2 µl in 18 µl MQ water (1:10) using template Pipetman (T2) different from the one used for primary PCR (can be multichannel or P20T2).

6. Rinse gloves in 0.1% bleach and 70% EtOH.

7. Open 2° PCR plate and add 1 µl diluted 1° PCR product to respective wells in 2° PCR plate . Close and seal 2° PCR plate and 1° PCR plate.

Run 2° PCR:

95 °C 2 minutes

| | | |
|---|---|---|
| 95 °C | 15 secs | 5 cycles |
| 60 °C | 30 secs | |
| 72 °C | 30 secs | |

| | | |
|---|---|---|
| 95 °C | 15 secs | 40 cycles |
| 55 °C | 30 secs | |
| 72 °C | 30 secs | |

72 °C 1 min
15 °C forever

Wipe Pipetman with 0.1% bleach and 70% EtOH.

Clear and seal waste and clean biosafety cabinet. Turn on biosafety cabinet UV and room UV for 30 minutes and then turn off.

**Gel electrophoresis**

Gels should be run in a different room from where PCRs are performed

Pour a 1.5% multi-tier agarose gel in TAE or TBE.
Open 2° PCR plate in gel room just before gel electrophoresis and not before.
Load 2° PCR product (15 ul) on gel along with NTC and 100 bp ladder using a separate P20 (P20Gel) or Multichannel (MGel) pipette.
Run gel. Photograph in UV Gel Doc system.

**Note:**

1. The inclusion of no template controls (NTC) in primary RT-PCR and secondary PCR (template for NTC in secondary PCR is product of NTC from primary RT-PCR) for every

round of testing is critical for reliable testing. A negative NTC is a prerequisite for interpreting the results.
